# Identification of molecular nociceptors in *Octopus vulgaris* through functional characterisation in *Caenorhabditis elegans*

**DOI:** 10.1101/2025.08.19.670888

**Authors:** Eleonora Maria Pieroni, Vincent O’Connor, Lindy Holden-Dye, Pamela Imperadore, Graziano Fiorito, James Dillon

## Abstract

Nociception, a phenomenon crucial for animal survival, deploys evolutionarily conserved molecular mechanisms. Among invertebrate species, cephalopods are of particular interest as they possess a well-developed brain speculated to be able to encode pain-like states. This has led to their inclusion in the Directive 2010/63 EU for welfare protection.

However, the molecular mechanisms of nociception in cephalopods are still poorly characterised and it is important to address this knowledge gap to better understand cephlapods’ capacity to express pain states. Here we describe a bioinformatic pipeline utilising conserved nociceptive genes, to identify the orthologous candidates in the *Octopus vulgaris* transcriptome. We identified 51 genes we predict to function in nociception. These add to the mechanosensory TRPN and the unique chemotactile receptors recently identified in octopus suckers, thus expanding the set of genes that merit further functional characterisation in cephalopods. We therefore selected 38 orthologues in *Caenorhabditis elegans,* a tractable experimental platform and tested loss of function mutant strains of distinct functional gene classes (e.g., *osm-9, egl-3, frpr-3*) in a low pH avoidance paradigm. This identified 18 nociceptive-related genes to be prioritised for further functional characterisation in *O. vulgaris*.

## Introduction

The ability to detect potential noxious stimuli in the environment involves the triggering of specialised sensory neurons called nociceptors which activate a simple reflex response that organises a withdrawal behaviour of the animal from a potential threat (Sherrington, 1906; Dubin and Patapoutian, 2010). The number and type of modalities of noxious stimuli detected by these cells is encoded by the molecular determinants which define the transduction and integration of such environmental cues (Wood, 2006; Dubin and Patapoutian, 2010). The crucial adaptive value of nociception in contributing to increase the individual survival, made it a conserved feature across Eumetazoa (Smith and Lewin, 2009).

In cephalopods, evidence of nociception is found in *ex vivo* and *in vivo* experimental observations in which noxious cues delivered to the arms or mantle of the animal, is able to trigger withdrawal responses and associated neurophysiological features such as post-injury sensitization (Hague et al., 2013; Crook et al., 2013; Alupay et al., 2014; Crook et al., 2014; Howard et al., 2019). This interpretation is reinforced in studies where a complex response is suggested to be organised in the central brain to produce secondary protective behaviours such as grooming and concealing of the arm after noxious exposure (Crook, 2021). The increasing interest in nociception and its processing in these cephalopods derives from the recognition of their sophisticated neuroanatomical central organization. This likely entails a top-down regulation of modulatory nerve signalling underlying the potential to exhibit pain pathways. Due to this possibility, as a precautionary principle, cephalopods are currently subject to legislation that protects their welfare when used in research (European Parliament and Council of the European Union, 2010; Smith et al., 2013; Birch et al., 2021). However, to fully address the question around pain perception, a broader understanding of the molecular determinants of nociception in cephalopods is required. To date, only two molecular classes of sensory detectors have been experimentally identified in *O. bimaculoides* (van Giesen et al., 2020) and *Sepioloidea lineolata* (Kang et al., 2023). These are located in the sensory epithelium surrounding the suckers which harbours putative cellular nociceptors that morphologically resemble mammalian sensory neurons (Rossi and Graziadei, 1956). One was identified as a NOPMC (TRPN) orthologue, activated by mechanical stimulation whilst the other class was found to be a phylum-specific ion channel, sensitive to both attractants and aversive cues (Kang et al., 2023). This provides evidence of both conserved and exclusive molecular components of sensory detection in cephalopod (van Giesen et al., 2020).

Most of the direct investigation of nociceptive processing in cephalopods is constrained by their limited experimental genetic tractability. These limitations derive from a series of challenges posed by culturing these animals, especially considering the very delicate early life stages (Iglesias et al., 2007; Vidal et al., 2014) and the strict requirements this species needs in terms of accommodation, nutrients, and temperature supplies (Ponte et al., 2022). Therefore, currently most of the work carried out on *O. vulgaris* is based on animals that are caught from the wild in a non-standardised way (Pieroni et al., 2022; Sykes et al., 2023). This generates two major issues, one is the complete absence of control over their genetic background and the second one is represented by the variable animal welfare status after capture and transport. However, the highly conserved genetic basis of nociception across the animal phyla provides an opportunity to take an indirect, but nonetheless, informative approach.

Here we conducted an *in silico* analysis to identify molecular candidates for nociception in the *O. vulgaris* transcriptome. To provide insight into their function we identified their *C. elegans* orthologues and tested loss of function mutants for deficits in a simple assay of chemical nociception by exposing mutant nematode strains to low pH, a common noxious cue in both species (Sambongi et al., 2000; Hague et al., 2013; Crook, 2021).

This approach highlighted 18 nociceptive-related genes prioritised for functional characterisation, reinforcing the molecular conservation of gene families in distinct organisms and mapping out a platform through which experimentally intractable cephalopod genes can be investigated in *C. elegans*.

## Materials and Methods

### In silico analysis

#### Resources to produce a list of conserved Eumetazoan nociceptive related genes

An in-depth literature review of conserved molecular nociceptors was performed by surveying around 400 peer reviewed journal articles on animal nociception from well-characterised vertebrate and invertebrate Eumetazoa.

This work was complemented with information taken from databases focussed on human pain genetics: 1) [Human Pain Genes Database](https://humanpaingeneticsdb.ca/) (Meloto et al., 2018) a collection of nociceptive relevant genes resulted from 294 studies reporting 434 associations between genetic variants and pain phenotypes; 2) [Pain Research Forum](https://painresearchforum.org/), curated by the International Association for the Study of Pain (IASP) and 3) [Pain Genetics MOGILAB](https://mogilab.ca/research/pain-genetics/) (McGill University, Montreal, Canada).

Finally, the queries “pain perception” and “opioid activity” were searched within Gene Ontology (GO) Resource (Carbon et al., 2009). The latter was included on the basis that this search term would highlight candidate molecular components that mediate or modulate anti-nociceptive pathways. The results under the “genes and gene products” label were selected for further analysis.

#### Gene blast against *O. vulgaris* transcriptome and manual curation of sequences

The search for orthologue nociceptive and antinociceptive candidate genes in *O. vulgaris* was performed by iterating the Uniprot amino acid sequence (The UniProt, 2025) against *O. vulgaris* transcriptome (Petrosino et al., 2015; Petrosino et al., 2022). A standard e-value of 10e^−5^ was used as a cut-off for the BLAST search except for neuropeptide precursors for which it was set as 10e^−2^. The rationale for the latter reduced stringency is due to the known propeptide variation relative to highly conserved and short length mature and active peptide component (Baggerman et al., 2005; Akhtar et al., 2011). When multiple hits were found, we considered the contig sequences with the e-value closest to 0 and a high query coverage (at least 40%) for further analysis. Incomplete sequences from the BLAST research that we failed to reconstruct were discarded. A set of bioinformatic tools was utilised to check the quality of the pre-existing automated annotations by manually curating the sequences. The nearest species with a sequence matching the candidate provided was retrieved from NCBI Blast (Altschul et al., 1990). A prediction of the protein function pathways (GO terms for “Biological Process”, “Molecular Function” and “Cellular Component”) was obtained via InterProScanSearch (Blum et al., 2021) following translation of the target transcript on Expasy Translate (Gasteiger et al., 2003). The analysis of the conserved domain, the family and shared structure of the protein was also performed, using NCBI conserved domain (Lu et al., 2020) and HHPRed (Hildebrand et al., 2009).

#### Refinement of candidate *O. vulgaris* nociceptive genes

Refinement of the final *O. vulgaris* sequences to be blasted against the *C. elegans* genome was performed by taking into account the available relative gene expression of the candidates in *O. vulgaris* (Petrosino et al., 2015; Petrosino et al., 2022). To this end, specific tissues were considered, based on their relevance in neurosensory pathways: tip of the arm (TIP), selected due to previous studies showing it is enriched in sensory receptors including those involved in nociceptive responses (Graziadei, 1964; van Giesen et al., 2020); supra-oesophageal mass (SEM), sub-oesophageal mass (SUB) and optic lobe (OL), constituting the central brain mass fundamental in information processing and thus potentially implicated in the elaboration of nociceptive and anti-nociceptive responses. This process led to the selection of a specific representative for each gene when multiple plausible hits were found.

#### Identification of *C. elegans* orthologues for candidate *O. vulgaris* nociceptive genes

*C. elegans* orthologues of the hits derived from the approaches described above, were retrieved from Wormbase resource (Davis et al., 2022) through the available BLAST tool (version WS288, Bioproject PRJNA13758).

To compare the degree of molecular conservation between *O. vulgaris* and *C. elegans*, the respective predicted protein translations were aligned using Clustal omega (Madeira et al., 2022) and the identity of the protein functional domains was compared with Expasy-SIM tool using BLOSUM62 as the comparison matrix (Huang and Miller, 1991). Where multiple hits were found, the selection was made based on literature evidence for their implication in sensory responses or on their expression in sensory neurons (e.g., npr-17 as allatostatin C receptor representative within its suggested 13 orthologues; Beets et al., 2023).

### *C. elegans* strains and husbandry

The following nematode strains were utilised in this study: CB1124 *che-3 (e1124);* CB75 *mec-2 (e75)* [STOM]; TU253 *mec-4 (u253*) [ASIC]; CB1611 *mec-4* (e1611) [ASIC]; CB1515 *mec-10 (e1515)* [ASIC]; *acd-1/deg-1 (bz90/u38u421)* [FaNaC]; TQ5 *trp-1 (sy690)* [TRPC]; RB1052 *trpa-1 (ok999)*[TRPA]; TQ6 *trp-4 (sy695)* [TRPN]; CX10 *osm-9 (ky10)* [TRPV]; CX4544 *ocr-2 (ak47)* [TRPV]; FG125 *ocr-2 (ak47), osm-9 (ky10), ocr-1 (ak46)* [TRPV]; EJ1158 *gon-2 (q388)* [TRPM2]; LH202 *gtl-2 (tm1463)* [TRPM3]; PT8 *pkd-2 (sy606)* [TRPP]; AG405 *pezo-1 (av143)* [PIEZO]; RB883 *kqt-2 (ok732)* [KCNQ]; TM542 *kqt-3 (tm542)* [KCNQ]; RB1392 *shk-1 (ok1581)* [KCNA]; *twk-46 (tm10925)* [KCNK]; CB251 *unc-36 (e251)* [CACNA2D3]; VC48 *kpc-1 (gk8)* [PCSK1]; MT150 *egl-3 (n150)* [PCSK2]; MT1218 *egl-3 (n588)* [PCSK2]; VC671 *egl-3 (ok979)* [PCSK2]; MT1071 *egl-21 (n476)* [CPE]*; acn-1 (tm12662)* [ACE]*; acn-1 (tm8421)* [ACE]; BR2815 *nep-1 (by159)* [NEP]; JN356 *nep-2 (pe356)* [NEP]; VC2171 *tkr-1 (ok2886)* [TKR]; RB1340 *nlp-1 (ok1469)* [TK]; VC2565 *frpr-3 (ok3302)* [FMRFaR]; NY7 *flp-1 (yn2)* [FMRFa]; NY16 *flp-1 (yn4)* [FMRFa]*; npr-17 (tm3210)* [AstCR/OPRL]; RB2030 *nlp-3 (ok2688)* [AST/OP-like]; FX03023 *nlp-3 (tm3023)* [AST/OP-like]*; anoh-1 (tm4762)* [ANO]; CX14295 *pdfr-1 (ok3425)* [CGRPR]; RB1546 *tmc-1 (ok1859)* [TMC]; FK100 *tax-2 (ks10)* [CNG]; PR671 *tax-2 (p671)* [CNG]; PR678 *tax-4 (p678)* [CNG]; BR5514 *tax-2/tax-4 (p671/p678)* [CNG]; VC9 *nca-2 (gk5)* [NALCN]. The Bristol N2 was used as the wild-type strain.

*C. elegans* were cultured and maintained as described in Brenner (1974). Single colonies of saturated LB broth cultures containing *Escherichia coli* OP50 were used to seed 15 mL nematode growth medium (NGM) agar plates (50 μL per 55 mm diameter plate). Plates containing *C. elegans* were then sealed with parafilm to prevent contamination and kept at 20°C. Three days prior to any assay, gravid adult worms were put on culturing plates to lay eggs for 4h and then removed to produce an age-synchronised population to be tested at the L4+1 day (young adult) stage.

### Drop assay to test acidic aversion in *C. elegans*

On the experimental day, ten L4+1 day old worms were transferred onto a 9 cm unseeded NGM plate (20mL) and left undisturbed for 20 min, in order to favour the transition from local area search (with high levels of reversal) to dispersal behaviour with low reversals and forward movements (Gray et al., 2005). A small drop of noxious cue was delivered in front of a moving worm through a small electrophysiology glass capillary (1.0mmOD, Harvard Apparatus, USA) attached to a 3 mL plastic syringe (Fisherbrand™). The number of reversals exhibited by the worm within 5s of exposure to the cue was recorded. A binary score was assigned with a positive response (1) scored for worms reaching the threshold displayed by the WT N2 (at least three complete reversals within 5s from the exposure to the substance), otherwise a negative response (0) was assigned. Worms for each strain for each condition were tested by exposing them to the drop only once. The resulting response score was calculated and compared to the WT N2 performance. Strains with known or observed strong impaired movements were not included in the study as they could have potentially affected the locomotory readout on which the assay is based.

To trigger low pH response, acetic acid (CAS No. 64-19-7, Fisher Chemicals™) was dissolved in M9 buffer to reach a final pH of 3 (M9, pH3). This cue was compared to the exposure of M9 buffer alone (M9, pH7). The experimenter was blind to the genotype being tested and exposure to cues was randomised. Data were analysed using either one-way or two-way parametric analysis of variance (ANOVA) with different substances (M9, pH3 vs M9 pH7, Na acetate vs M9) and strains vs WT N2 as between-subjects.

Post-hoc comparisons were performed using Dunnett’s multiple comparisons test for one-way ANOVA and Tukey’s multiple comparisons test for the two-way ANOVA. A level of probability set at p<0.05 was used as statistically significant. Statistics were performed with GraphPad Prism version 10 for Windows (GraphPad Software, Boston, Massachusetts, USA).

## Results

### Resolution of a query set of conserved Eumetazoan nociceptive-related genes

Our bioinformatic pipeline, identified a total number of 474 nociceptive related genes collectively retrieved from literature analysis (141), databases (200) and GO search (133).

A considerable number of retrieved genes (152 entries) was found multiple times across the different sources or under synonyms. These duplications were removed, resulting in a total number of 322 distinct candidates (Fig. 1A and Table S1A).

**Fig. 1.**
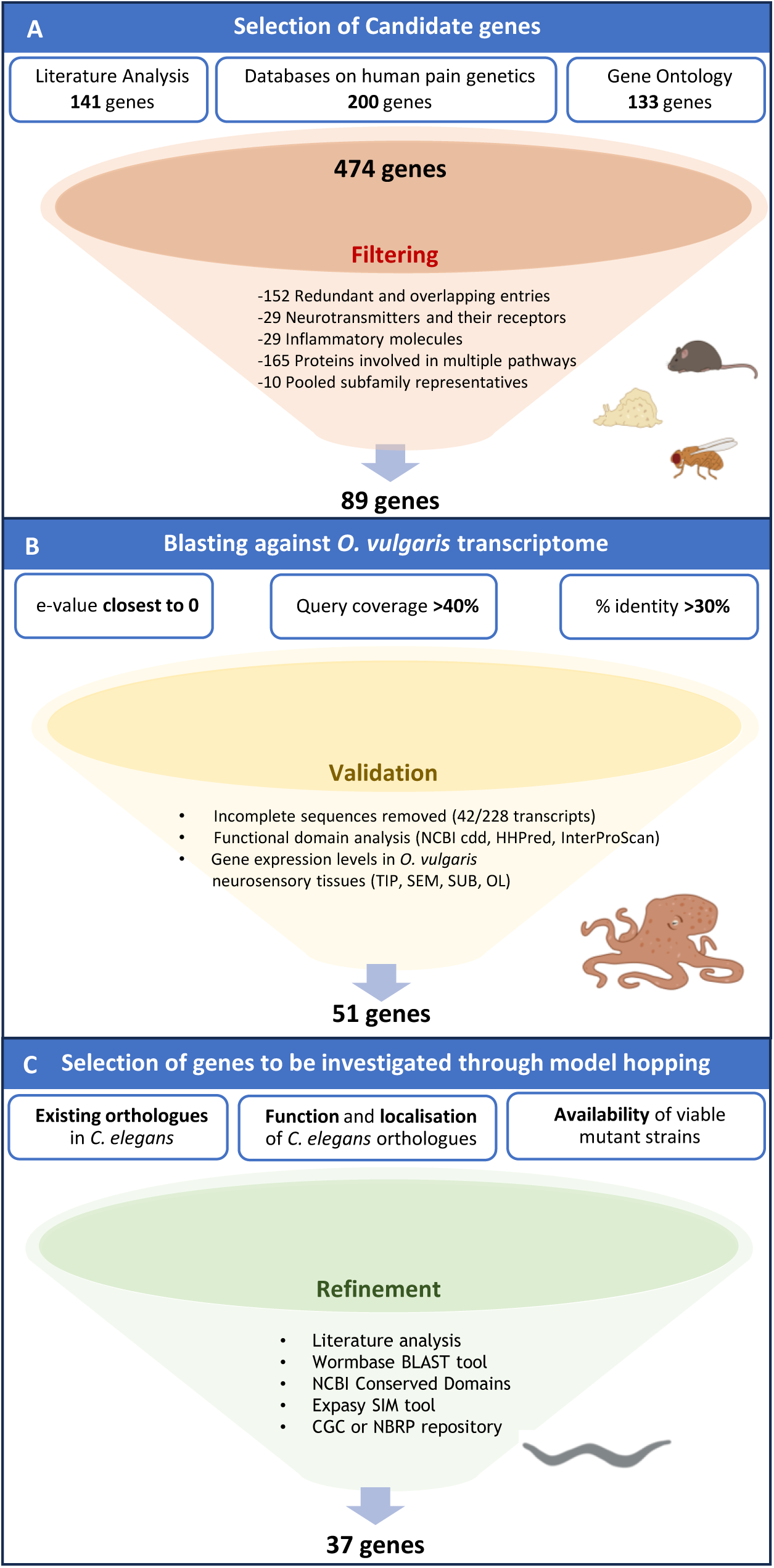
Summary of *in silico* analysis leading to the selection of *O. vulgaris* putative molecular nociceptive determinants to be modelled in *C. elegans*. **(A)** Literature analysis of well-characterised Eumetazoan nociceptors was complemented with information from databases on human pain genetics (Human Pain Genetic Database, IASP Pain research, MOGILab) and GO “gene and gene products” to produce 474 distinct candidate genes. We manually filtered to exclude overlapping entries among resources (152), genes involved in multiple pathways (165), neurotransmitters and their receptors (29) and those involved in the inflammatory component of pain (29). We additionally pooled together (10) different representatives of the same subfamily (e.g., ASIC1-3, TRPV1-8 etc.) **(B)** The resulting 89 candidates were blasted against *O. vulgaris* transcriptome to retrieve the closest orthologues which were then manually curated, leading to 51 candidate genes. We prioritised genes based on available tissue expression levels in octopus sensory and nervous tissues (TIP, SEM, SUB, OL – for details see Table S2). **(C)** To promote the functional characterisation in *C. elegans,* mutant strains for the orthologue genes were selected This filtering led to a final number of 37 genes across 45 strains.

We then filtered down this list by focusing on essential nociceptive and anti-nociceptive determinants, such as receptors directly gated by noxious stimuli or known modulators of sensory detection. This was done with a view of excluding categories of proteins involved in multiple physiological pathways or with a rather secondary/diffuse role in nociception. Among these, neurotransmitters, and their receptors (29 genes), transcription factors, large families of enzymes involved in widespread cellular activities (165 genes) and molecules contributing to the inflammatory responses were excluded (29 genes). Furthermore, given the challenge posed by some candidates in distinguishing between isoforms of the same gene or different representatives of the same subfamily we pooled related genes (e.g., TRPV1-6 have been pooled into TRPVs, ASIC1-3, into ASICs, PIEZO1-2 into PIEZO) for a total of 10 genes.

Altogether, this filtering produced a final number of 89 potential candidate genes that were blasted against *O. vulgaris* transcriptome (Fig. 1A and Table S1B).

### *O. vulgaris* shares most of the conserved protein families implicated in nociception

Each selected protein sequence from the query set was blasted against *O. vulgaris* transcriptome to find orthologue genes (Fig. 1B).

The search of the 89 genes in *O. vulgaris* transcriptome led to a total number of 228 transcripts. We then manually curated each of them to remove incomplete sequences, identical sequences, and we managed to reconstruct two transcripts from fragments. This refinement led to a total number of 171 complete transcript sequences (Table S2). The corresponding 51 distinct genes encoded for proteins belonging to more than 30 different families (and 50 subfamilies) of receptors and modulators associated with sensory detection of chemical, mechanical and/or thermal stimuli. The results were organised into different categories based on the putative biological role of these proteins: 1) Classical activators of nociception, 2) Voltage-gated ion channels/subunits, 3) Proteins related to neuropeptide and lipid metabolism, 4) Other modulators of nociception (Table S2).

Belonging to the first category, orthologues of well-characterised proteins such as transient receptor potential (TRPs) and amiloride sensitive (ASCs) ion channels were identified with 9 and 3 representatives (within multiple isoforms) respectively.

The presence of specific subunits of ion channels implicated in nociception, such as the calcium voltage-gated channel subunit α2δ subunit 3 (CACNA2D3), known for its role in thermosensation was found, whilst other known channels specifically involved in nociception triggering such as Na_v_1.7-Na_v_1.9, were not clearly identified, mostly leading to Na_v_1.1 (SCNA1) and Ca_v_1.1 (CACNA1S) orthologues (Table S2).

### Genes implicated in canonical analgesic pathways are missing from *O. vulgaris* transcriptome

Out of the 89 selected genes, 38 were not found in *O. vulgaris* transcriptome. These include opioid precursors and derivatives, classical opioid receptors, and endocannabinoid receptors. However, receptors that share strongest homologies with those receptors, such as Allatostatin C receptor (AstCR) with an assigned “opioid receptor” protein family membership (IPR001418) in InterProScan, and an Opioid growth factor receptor (OGFR) were identified. Furthermore, enzymes such as neprilysin (NEP), carboxypeptidase (CPE), neuroendocrine convertase (PCSKs) in the case of neuropeptides-related proteins or diacylglycerol lipase (DAGL) and N-acyl-phosphatidylethanolamine-hydrolysing phospholipase D (NAPE-PLD) in the case of lipid modulators were found. This shows the existence of conserved pathway for the biosynthesis, processing, and metabolism of neuropeptides and lipids that have a broader physiological role that may include nociception modulation and its physiological mitigation.

### *C. elegans* mutants of *O. vulgaris* orthologue genes were selected for characterisation in a chemical aversion assay

Following the identification and manual curation of *O. vulgaris* candidates, we looked at the data on available tissue distribution and prioritised the analysis of genes showed to be enriched in neurosensory tissues (TIP, SEM, SUB and OL) especially when more than one hit resulted in a plausible orthologue candidate of the putative nociceptive gene (Table S3).

As we intended to model the response triggered by these genes in *C. elegans*, each resulting *O. vulgaris* protein sequence was blasted against *C. elegans* genome database to find, where present (Table S3), the closest nematode orthologue (Fig. 1C). The result of this final refinement led to 37 *C. elegans* genes, for which we selected 45 different strains (Table 1).

**Table 1.**
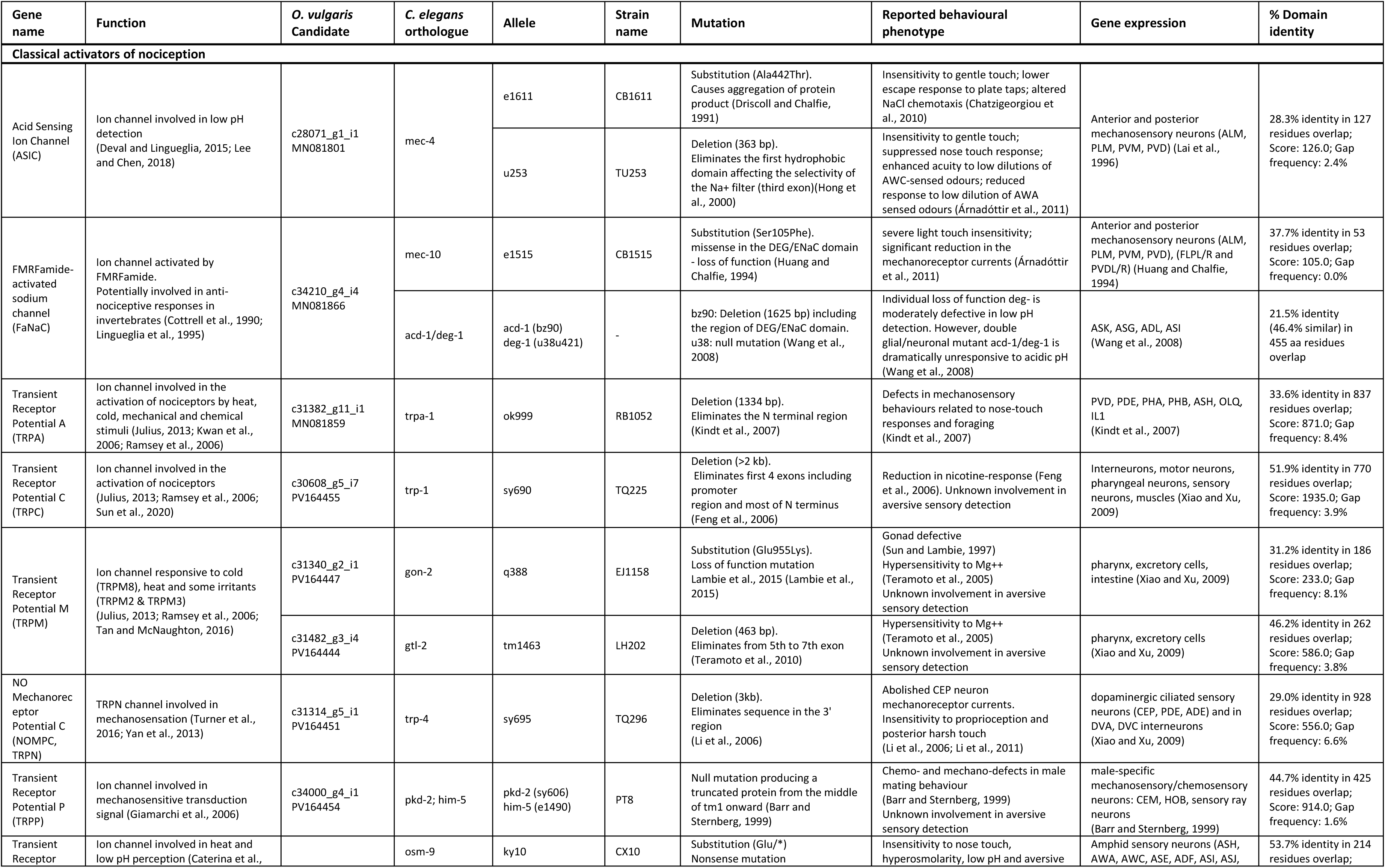
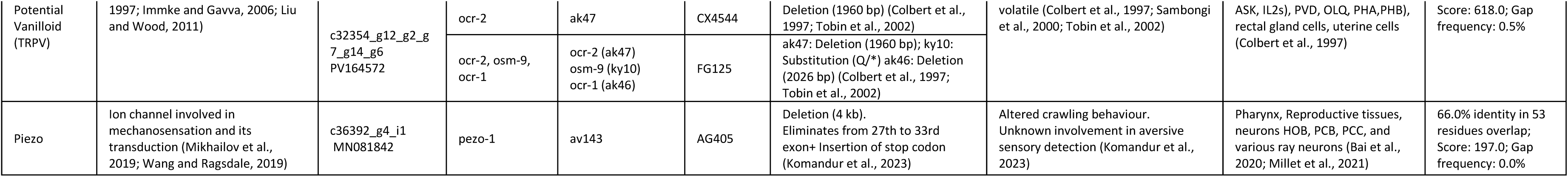
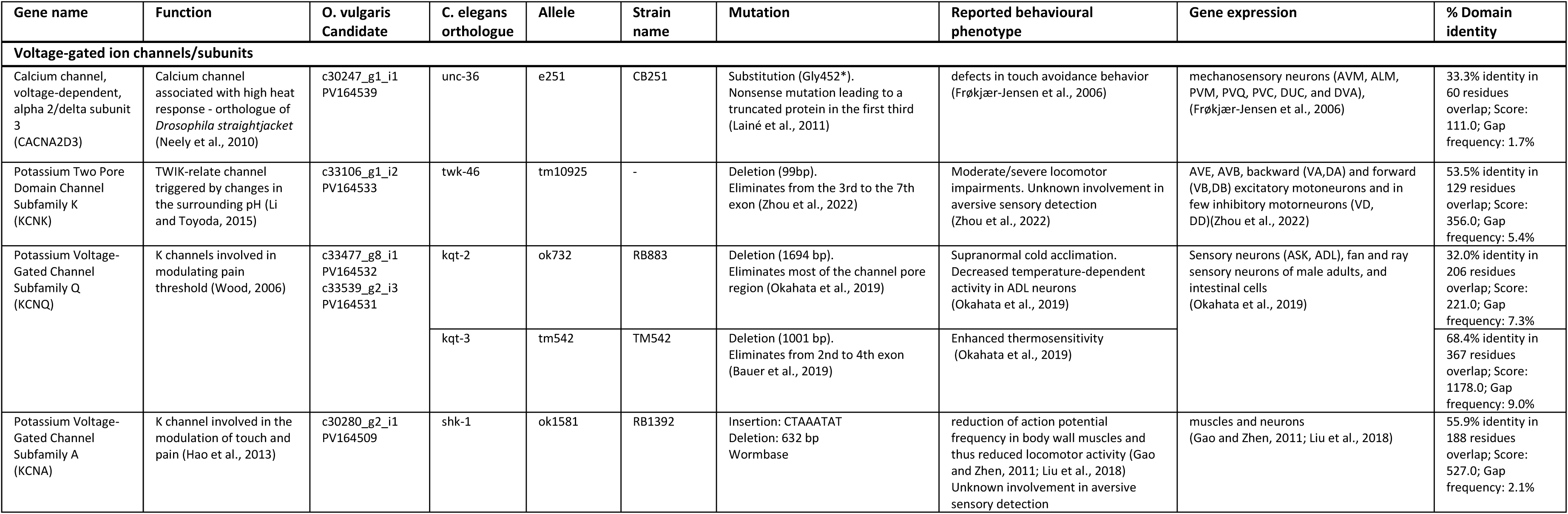
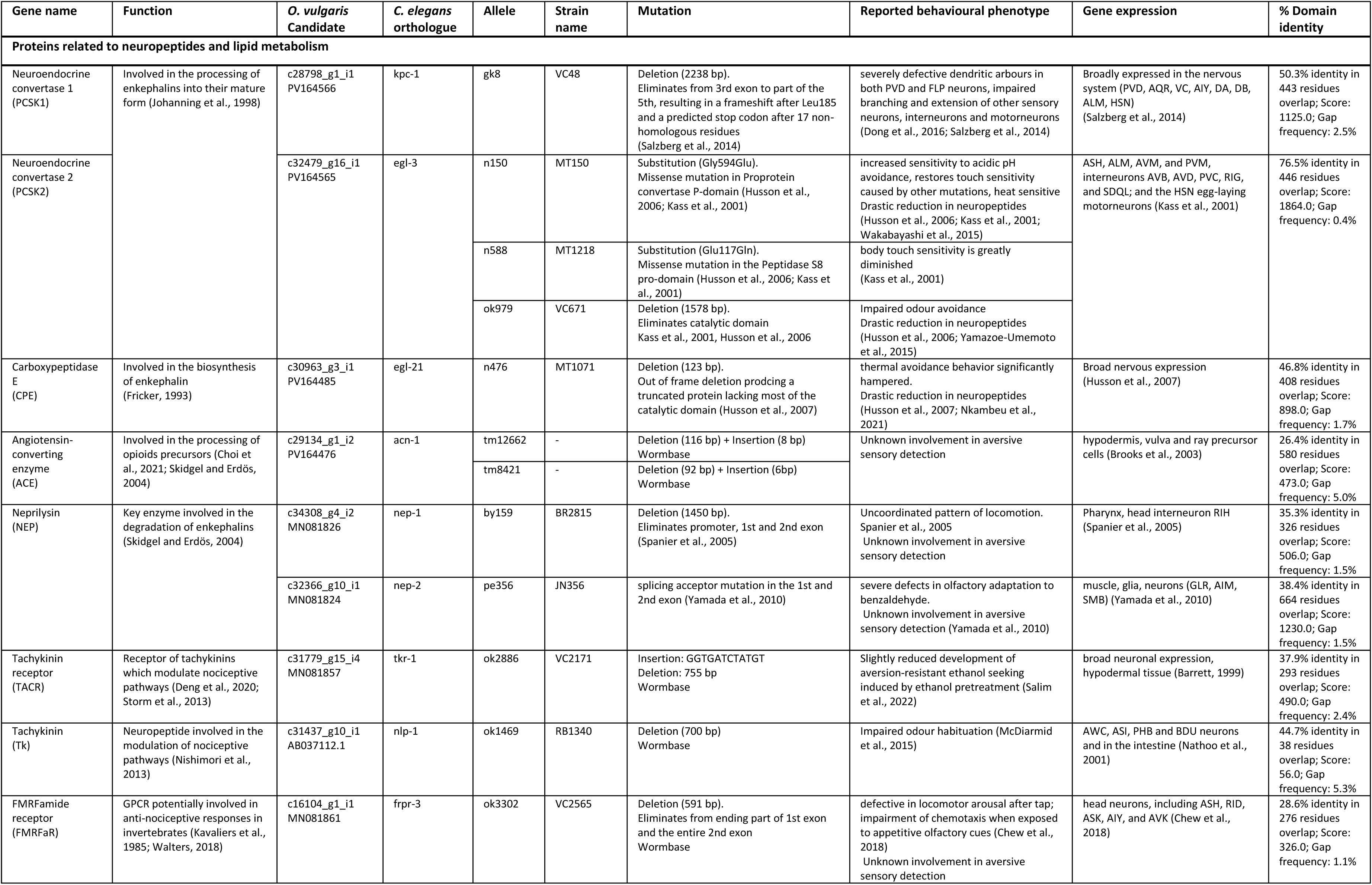
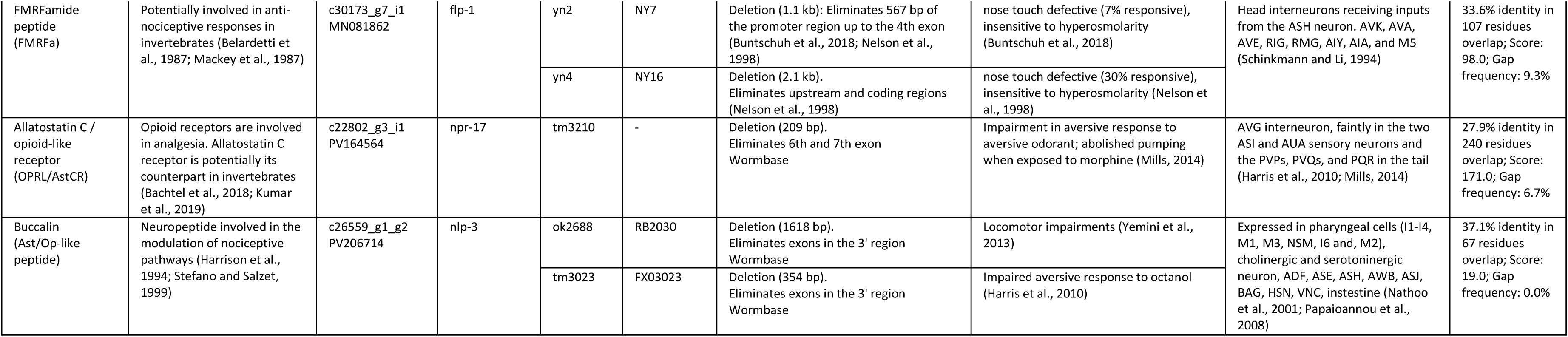
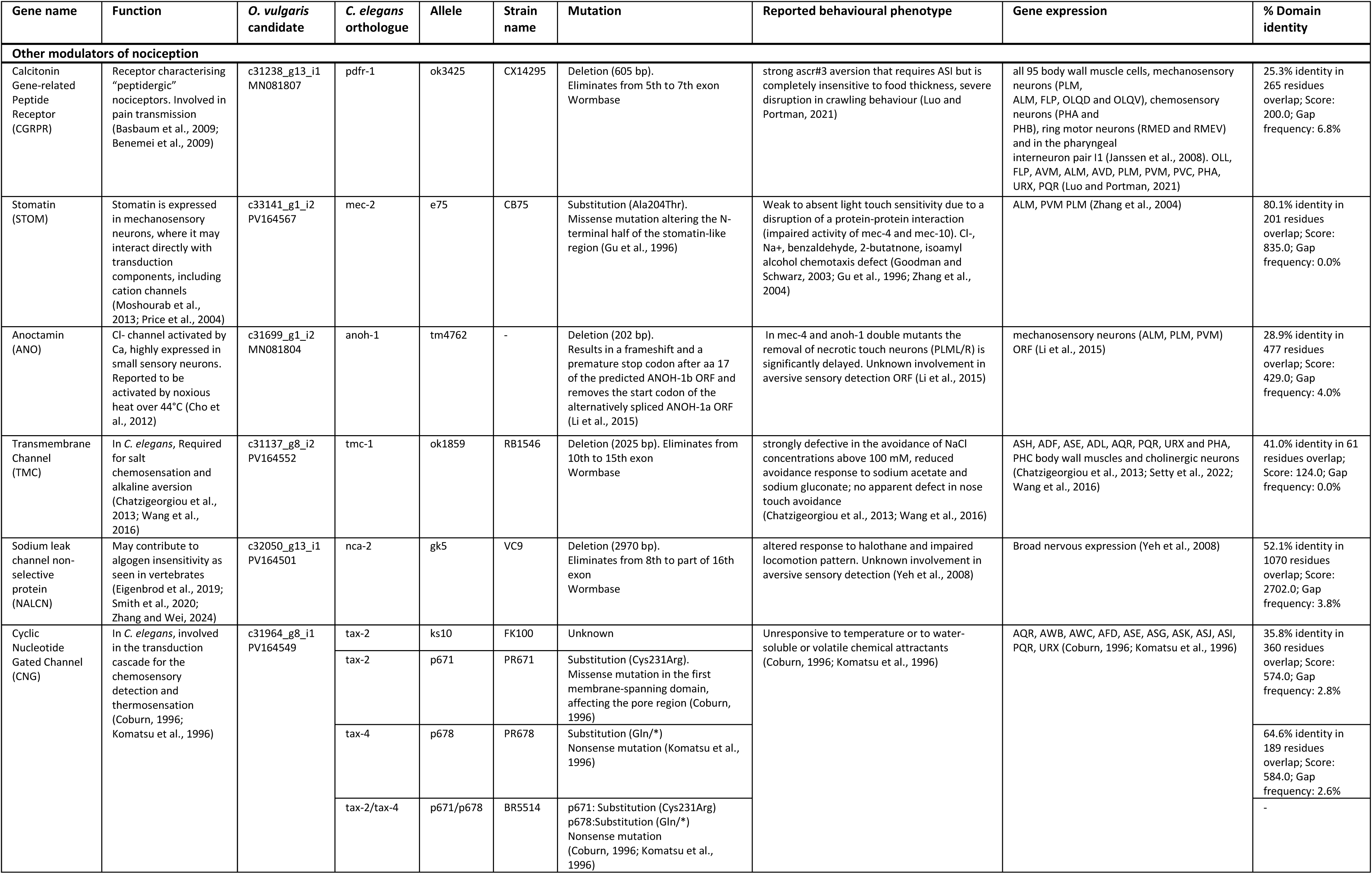
List of *C. elegans* loss of function mutant strains for the selected nociceptive related genes selected from *O. vulgaris* transcriptome after manual filtration of the *in silico* analysis. For each gene, the first four columns report the gene proposed function based on literature, the *O. vulgaris* candidate gene (based on the assembled transcriptome Petrosino et al., 2015 and Petrosino et al., 2022) and assigned NCBI Accession number. Columns 5-10 refer to the orthologue gene of *C. elegans*, the strain and alleles selected to be tested for chemoaversion impairments and the pre-existing knowledge on the behavioural phenotype of the mutant, as well as the known localisation of the gene in worms tissues. In some cases, a mutation in more than one allele encompassing the same gene was tested. The last column refers to the degree of identity and similarity found between the two species genes when comparing their functional domains (Expasy-SIM tool, BLOSUM62 comparison matrix).

Based on the recurring evidence showing nociceptive responses triggered in octopuses through acetic acid administration (*O. vulgaris*, Hague et al. 2013, *O. bocki,* Crook, 2021), we selected low pH (M9, pH 3) as the exemplar cue and systematically tested against selected *C. elegans* strains using the chemoaversion drop assay.

### The drop assay is a sensitive test to study chemoaversion in *C. elegans*

We analysed WT N2 and selected mutants in the assay and benchmarked the results against previously published data to validate our approach. The results, in agreement with published data (Sambongi et al., 2000) and showed that low pH elicited a significant aversive response in N2 (M9, pH3 average number of reversals = 3.990 ± 0.115) when compared to the control (M9, pH 7, average number of reversals = 0.432 ± 0.033). This allowed us to set the threshold for measuring responsiveness (1 for avoidance, 0 for defective avoidance) to at least 3 reversals for M9 pH3 (Fig. S1).

We then verified the sensitivity of the drop assay in discriminating the responsiveness of different strains of worms. We first tested the well-characterised *che-3 (e1124),* a strain with ultrastructural defects in the ciliated dendritic endings of the amphid sensory neurons that play an essential role in chemosensation and compared its performance to the WT N2 (Perkins et al., 1986; Wicks et al., 2000).

As previously published for *che* mutant strains (Sambongi et al., 2000), *che-3 (e1124)* showed a poor avoidance performance to acidic pH when compared to the WT N2 (p<0.0001) with less than 30% responsiveness to low pH (M9, pH 3 average number of reversals 2.293 ± 0.130), a difference of more than 50% with the WT N2 (around 80% responsiveness Fig. S1).

Additionally, we analysed *C. elegans* responsiveness to 200mM Na acetate (C₂H₃NaO₂) to check which chemical component of acetic acid was responsible for triggering the aversive response. Worms showed no sensitivity to sodium acetate (200 mM Na acetate vs Ctrl M9 pH7, p = 0.9802 in WT N2, p = 0.9924 in *che-3 (e1124)*) demonstrating that the repulsive response of M9, pH3 was elicited by the protons (H^+^) rather than by the acetate ion (Fig. S1).

### *C. elegans* is a discriminating platform for characterising conserved molecular nociceptors

Once we established the drop assay as a valuable test to discriminate the responsiveness of the strains, we analysed the performance of all the 45 strains of *C. elegans* mutants that came out of the list of 37 orthologues of *O. vulgaris* putative nociceptive genes (Fig. 2). The drop assay identified 18 distinct genes across the four different categories that seem implicated in the detection or processing of acidic pH.

**Fig. 2.**
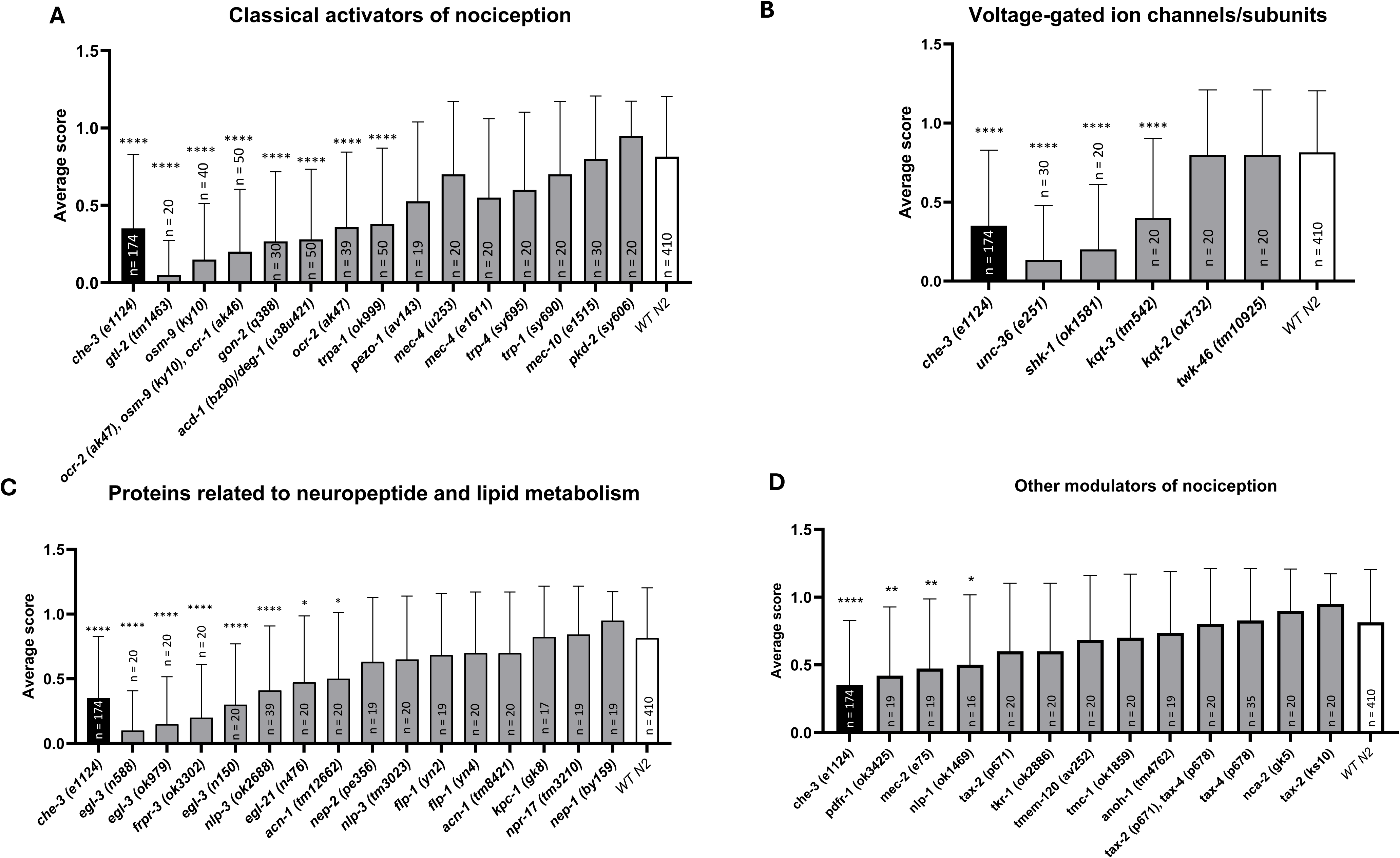
Functional investigation of *C. elegans* orthologues of *O. vulgaris* genes implicated in nociception. The bars represent the average score performed by the total number of worms per strain tested ± SD. The putative genes involved in nociception were grouped into distinct categories (A-D) and the acidic aversion of mutant strains for each gene was compared to the controls, WT N2 and *che-3 (e1124*). In each experiment, worms were exposed to a drop of acetic acid (pH 3) only once and the response was recorded and classified according to the designated criteria described in the text. Data were subjected to one-way ANOVA followed by Dunnett’s multiple comparisons test. Category A: “Classical activators of nociception”, refers to all the genes that were found recurringly across all the resources we interrogated (e.g., TRPs, ASCs, PIEZOs), Category B: “Ion channel and subunits involved in nociception” is self-explanatory, Category C “Proteins related to neuropeptide and lipid metabolism” was originally meant to include representatives from the opioid and endocannabinoid systems but given we couldn’t find any obvious orthologues, we focused on the enzymes involved in the synthesis, maturation or catabolism or neuropeptides and lipids that might still have a fundamental role for counteracting nociception. Category D: “Other modulators of nociception” refers to a broader range of genes that encode for enzymes and receptors that have been known to be implicated in nociception processing. *p<0.05, **p<0.01, ***p<0.001, **** p<0.0001 refers to the comparison against the WT N2 performance; n represents the number of individuals tested. Each strain was tested in at least two independent experiments.

Seven out of 18 genes belong to the “classical activators of nociception” category with impaired strains, all showing an average number of reversals below the set threshold for responsiveness (Fig. 3).

**Fig. 3.**
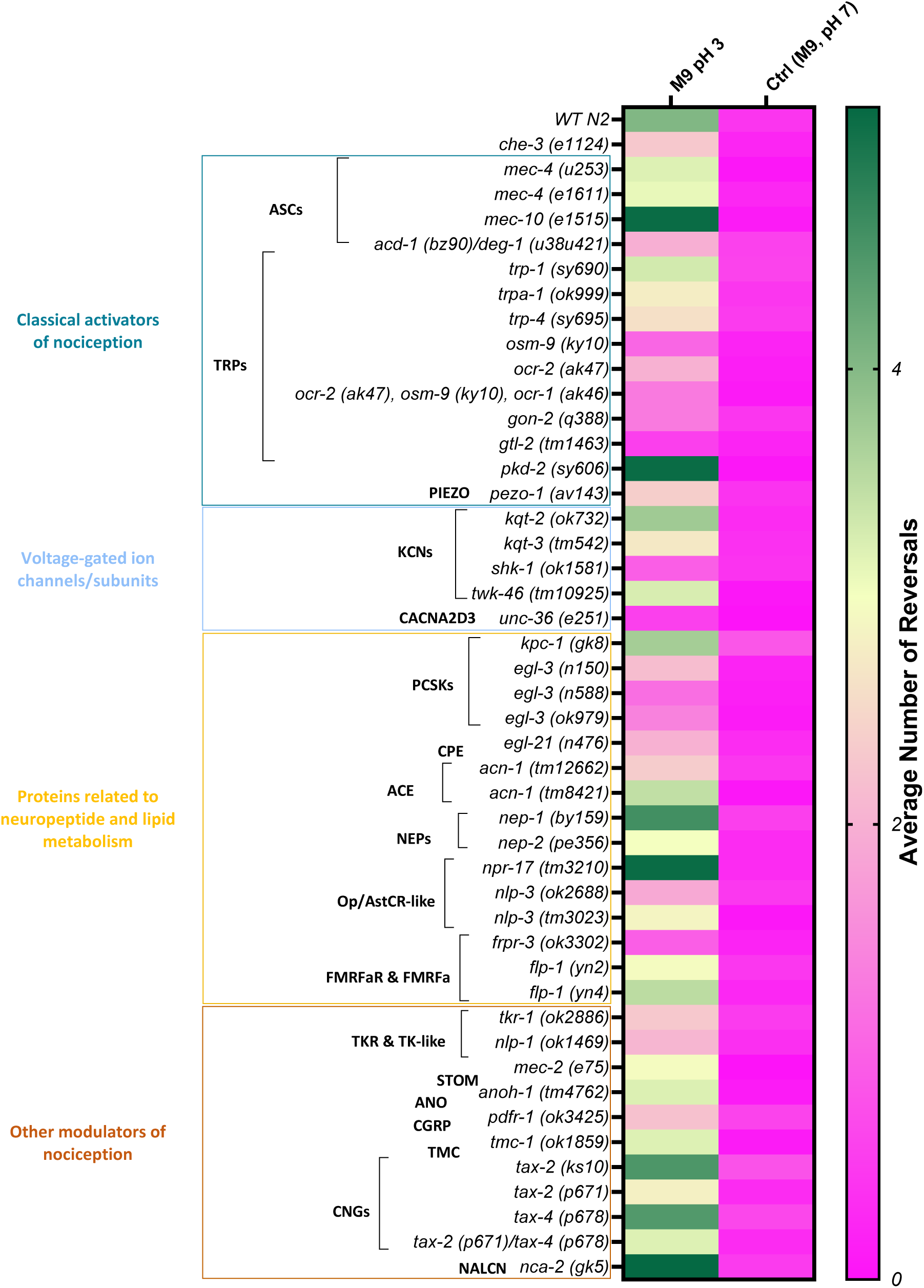
*C. elegans* highlights potential molecular determinants involved in *O. vulgaris* nociception. Data for the drop assay is reported here as the number of actual reversals after exposure to the noxious cue (M9, pH 3). The heat map shows a colour coded indication of the strains’ sensitivity to M9, pH 3 from non-sensitive *che-3 (e1124)*-like (magenta) to sensitive WT N2-like (green). Each colour block represents the average number of reversals across all the worms tested per strain. The WT N2 performance was characterised by at least 3 body reversals within 5s of contact with the drop and was used to set our criteria for responsiveness.

Mutant strains belonging to the TRP channel subfamilies, including the TRPVs *osm-9* (p<0.0001) and *ocr-2* (p<0.0001), the TRPA *trpa-1* (p<0.0001) and the TRPMs *gon-2* (p<0.0001) and *gtl-2* (p<0.0001), all showed a significant impairment in the detection of acidic pH (Fig. 2A). Representatives of TRPN (*trp-4*), TRPC (*trp-1*) and TRPP (*pkd-2*) TRP subfamilies, however, did not show any defect in pH 3 aversion.

Worm strains of the orthologues of other families of classical pH detectors such as ASCs, did not show any defect in chemoaversion as it might be expected by acidic sensing ion channel representatives. In fact, *mec-4* and *mec-10* strains exhibited on average more than three reversals (3.2000 *mec-4 (u253)*, 3.1000 *mec-4 (e1611)*, 5.1000 *mec-10 (e1515)*, Fig. 3). The only strain showing lack of response to pH 3, was the neuronal and glial ASC double mutant *acd-1 (bz90)/deg-1(u38u421)* (p<0.0001).

An impaired response to pH3 was also found among specific strains representative of the “voltage-gated ion channels/subunits” category. This included *kqt-3* (p<0.0001) and *shk-1* (p<0.0001) representatives of the KCNQ and KCNA sub families of potassium channels. This impact was selective as neither *kqt-2* (p = 0.9118)*twk-46* (p>0.9999 with an average of 3.2500 reversals, Fig. 3) were altered in their withdrawal response. *unc-36* (p<0.0001), orthologue of the CACNA2D3 subunit, showed a defective response when challenged with the noxious pH3 cue (Fig. 2B).

Strains with mutations in genes that encode for “proteins related to neuropeptide and lipid metabolism”, showed defects in the detection of acetic acid (Fig. 2C). The *nlp-3 (ok2688)* strain showed a clear impairment in response to the repellent (p<0.0001), whilst mutant strains of FMRFamide-like peptide orthologues *(flp-1)* were responsive. Among the FMRFamide receptors *frpr-3*, showed a loss of response (p<0.0001, average number of reversals 0.9500, Fig. 3). Considering *C. elegans* orthologues of the enzymes involved in the biosynthesis *(egl-21)*, maturation *(kpc-1, egl-3, acn-1)* and degradation *(nep-1, nep-2)* of neuropeptides, the PCSK2 *egl-3* representatives (p<0.0001), the CPE *egl-21* (p = 0.0109) and the ACE *acn-1* (p = 0.0209) mutant strains showed a reduced response when exposed to the pH3 repellent (Fig. 2C).

Belonging to a broader category of “other nociception modulators”, only mutants in the peptide *nlp-1*, the stomatin-like protein *mec-2* and the calcitonin gene-related peptide receptor (CGRP) *pdfr-1,* seemed to be involved in pH 3 avoidance (p = 0.0427, p = 0.076 and p = 0.010 respectively). Finally, most of the genes in this category did not show defects in this specific aversive response. This includes CNG representatives *tax-2* and *tax-4*, the anoctamin-1 orthologue *anoh-1*, all eliciting a WT-like performance (> 3 reversals) when exposed to acetic acid (Fig. 2D and Fig.3).

## Discussion

### Core conserved molecular nociceptors in *O. vulgaris* can be identified using functional characterization in *C. elegans*

The *in silico* pipeline revealed that a large number of genes showing conservation to known nociceptive genes in other species are present in *O. vulgaris.* In particular, at least two representatives were found to belong to the acid sensing ion channels (ASICs), including the classical molluscan FMRFamide-gated Na^+^ channel (Cottrell et al., 1983;Cottrell et al., 1990; Cottrell et al., 1997), two genes belonging to the PIEZO family, whilst for TRPs, 9 members (with multiple isoforms) were identified (Table S2). These include two TRPA1 channels, two TRPCs, two TRPMs, one TRPP, one TRPN and one TRPV channel, showing less variety of members when compared to vertebrates (Peng et al., 2015). This suggests either the vertebrate TRPs underwent specialisation during evolution (e.g., gene duplication and divergence) or that the fewer invertebrate representatives play multiple physiological functions as in the original ancestral gene (Valencia-Montoya et al., 2024).

When tested in *C. elegans*, one of the strongest behavioural impairments was found in mutant strains for the TRPV channel representatives *osm-9* and *ocr-2.* (Fig. 2A). These results are in line with previous studies showing their role in acidic pH sensation in *C. elegans* (Sambongi et al., 2000) reinforced by the essential role that orthologues play in vertebrate noxious responses (Dhaka et al., 2009).

Interestingly, other TRP channel orthologues were defective in chemoaversion implicating them in the acidic response. These include *trpa-1* and the TRPM representatives *gon-2* and *gtl-2*. The former has been previously found to be implicated in mechanical and thermal responses in *C. elegans* whilst a WT response was elicited when exposed to ASH-dependent noxious chemical cues tested (i.e. copper chloride, 1M glycerol, denatonium; Kindt et al., 2007). Our results could therefore depend on the complexity of the low pH response circuitry, which does not exclusively rely on ASH neurons but on a series of amphids response (Sambongi et al., 2000; Bergamasco and Bazzicalupo, 2006). To the best of our knowledge, the TRPMs have never been tested for chemical sensitivity but have been mostly studied for their role in reproduction (Sun and Lambie, 1997; Teramoto et al., 2010). Despite the low expression levels of *gon-2* in amphids and the selective expression of *gtl-2* in pharynx and excretory cells, the phenotype observed might indicate some indirect effect of these TRP proteins in the chemosensation executed by exposure to low pH.

The only ASC strain to show defects when exposed to acidic pH was the double mutant *acd-1/deg-1* which carries mutation in both neuronal and glial ASC proteins, suggesting a glial role in the processing of low pH in *C. elegans*, similarly to what previously reported in vertebrates (Chesler, 2003; Erlichman et al., 2004; Wang et al., 2008; Gourine and Dale, 2022). The octopus arm, the main structure involved in sensory interaction with environmental cues, includes different types of neurons and glial-like cells that might highlight the potential role of this cell type in acidic nociception as well (Young, 1971; Winters-Bostwick et al., 2024). The lack of impairment in the rest of the ASC protein family members tested here is in line with previous studies showing a main role in mechanical detection (Suzuki et al., 2003; Zhang et al., 2004; Árnadóttir et al., 2011; Schafer, 2015; Al-Sheikh and Kang, 2020).

### *O. vulgaris* lacks canonical vertebrate opioid signalling

According to Bateson’s criteria (1991) for eligibility to feel pain, animals must be endowed not just with nociceptors connected to a central brain mass but they should also possess a system able to counteract the inner disruption of the organism homeostasis, namely analgesic pathways. Therefore, we included candidates for opioid and cannabinoid receptors, and their ligand precursors. Blasting of ‘classical’ vertebrate opioid receptors (i.e. mu-, kappa-, delta-) produced two matches. The first one was the opioid growth factor receptor (OGFR), which has no structural resemblance to the classical opioid receptors and constitutes a different protein family (Zagon et al., 2002). The second and closest homology was to the allatostatin C receptor (AstCR), the protostome orthologue of the vertebrate somatostatin receptor (Koch et al., 2022). Opioid receptors are thought to have evolved from a duplication of the AstCRs (Thiel et al., 2021) and a few studies have reported a role of AstC in nociception and immunity in invertebrates (Bachtel et al., 2018; Li et al., 2021), in addition to growth and feeding regulation (Mizoguchi, 2016; Stay and Tobe, 2007). This raises the intriguing possibility that AstC signalling might underpin anti-nociceptive signalling in phyla that lack an opioid system. In line with this, we selected the opioid-like/allatostatin C-like receptor *npr-17* as the best candidate to be tested in *C. elegans* (Beets et al., 2023). Previous work has shown that the receptor NPR-17 and the neuropeptides NLP-3 and NLP-24 are essential in the morphine-mediated 1-octanol avoidance response and that the impaired response of the *npr-17* null mutant could be rescued by the human κ-opioid receptor (Mills et al., 2016). However, in our experiment, mutant strains for all three genes were not impaired in low pH chemoaversion, confirming a more complex peptidergic modulation of the aversive response. A recent large-scale de-orphanisation of *C. elegans* GPCR identified an additional 12 AstCRs which might be worth analysing in the appropriate noxious context (Beets et al., 2023).

*C. elegans* orthologues of enzymes involved in the biosynthesis (i.e., the CPE *egl-21)*, and processing of neuropeptides (i.e., the PCSK2 *egl-3)*, showed disrupted responses to the low pH cue. Mutations in *egl-3* and *egl*-*21* were already known to show defects in mechanosensory and thermosensory avoidance responses (Kass et al., 2001; Nkambeu et al., 2019). Such defects can be related to the disrupted production of neuropeptides as highlighted by mass spectrometry experiments (Husson et al., 2006; Husson et al., 2007). Collectively, these data point out the importance of neuropeptide signaling in the processing of nociception (Dickinson and Fleetwood-Walker, 1998) but do not constitute evidence of the presence of the classical analgesic pathways. Indeed, even in the case of the opioid sensitive analgesic pathway there is a broader functional consequence of this signaling extending to underpinnings of appetite, gut function and systemic homeostasis (Yuan et al., 2012).

Our *in silico* findings are somewhat in line with previous phylogenetic and bioinformatic analysis suggesting opioids and endocannabinoids as vertebrate exclusive proteins (Thiel et al., 2021). This does not necessarily mean that invertebrates do not possess a system to counteract noxious stimuli as they could have more ancient or unique solutions to the same problem. FMRFamide has been considered the equivalent of opioid molecules in invertebrates as the sequence (Phe-Met-Arg-Phe-NH2) shows some similarities to the heptapeptide Met-enkephalin (Tyr-Gly-Gly-Phe-Met-Arg-Phe). Among the wider physiological functions these peptides were found to modulate in molluscs and more generally in invertebrates, there are a few showing an induced suppression of primary nociceptors in *Aplysia* (Belardetti et al., 1987; Mackey et al., 1987).

The mutant of the *C. elegans* FMRFa orthologue we selected, *flp-1* did not show impairment in acidic response but has been found involved in high osmolarity aversion (Nelson et al., 1998; Li et al., 1999) suggesting a more selective chemosensory role. The orthologue of the FMRFamide receptor, *frpr-3* on the other hand, showed a strong impairment in pH 3 response, and thus represents a strong candidate for further analysis.

### Lipid signalling is an important component of nociception modulation across phyla

In the case of the endocannabinoid pathway, our *in silico* analysis identified different *O. vulgaris* genes encoding proteins implicated in fatty acid metabolism, such as DAGL, NAPE-PLD or Fatty acid-binding proteins (FABPs) (Table S2). However, we did not find molecular homologies to sign post any classical receptors of these signaling cascades within the octopus genomes. Although we found no evidence for the canonical endocannabinoid receptors in octopus, the lipids that act on them have established physiological effects on other proteins such as allatostatin receptors and TRPVs.

Interestingly, one of the proposed “ancestors” of the endocannabinoid system is the TRPV channel, previously shown to mediate reduced nocifensive response when activated by anandamide and 2-acyl-gycerol in the medicinal leech *Hirudo verbana* (Summers et al., 2017). Specifically in the case of *C. elegans*, bioinformatic analysis coupled with thermal proteomic profile, identified the AstCR *npr-32*, and the TRPV *ocr-2* as responsible for modulating the nocifensive response in a thermal avoidance assay, confirming these molecular determinants as key players in the detection and modulation of nociceptive responses (Abdollahi et al., 2024).

### Limitations and potential confound of the study

Our *in silico* approach was biased towards the selection of pre-existing, conserved molecular determinants, thus excluding potentially phylum-specific proteins as recently demonstrated by the identified chemotactile receptors in octopus and squid (van Giesen et al., 2020; Kang et al., 2023). Our study found a large number of conserved candidates in the transcriptome of *O. vulgaris* (Petrosino et al., 2015; Petrosino et al., 2022). However, the database available is not refined and therefore manual curation led to either the exclusion of incomplete or misannotated transcripts or required manual reconstruction (e.g., TRPV) to be fully investigated. The availability of the recently published chromosome-level genome could potentially improve the quality of this pipeline (Destanović et al., 2023).

Some limitations in the translation of our results to *C. elegans* warrant discussion. In fact, a few *O. vulgaris* gene families representative of nociception signaling could not be modelled in the nematode due to the absence of corresponding orthologues. This is exemplified by the purinergic receptors (P2X) and voltage gated sodium (Na_v_s) channels (Bargmann, 1998; Harte and Ouzounis, 2002; Hobert, 2005-2018; Burnstock and Verkhratsky, 2009; Table S3). Furthermore, both the phylogenetic distance and the distinct environments in which the two species evolved should be considered, as the diverse modalities and variety of ecological stimuli could have driven shared proteins to diverge and acquire different adaptations (van Giesen et al., 2020; Allard et al., 2023).

### Future directions

By performing a chemosensory assay on *C. elegans* mutants for the orthologue of *O. vulgaris* putative nociceptive genes, we have highlighted candidates that are involved in the acidic pH avoidance response. Two main categories of proteins deserve priority for further investigation: One is the classical activators of nociception, such as the TRPV channel subfamily, whose role in nociception is widely conserved across phyla, and the other is the neuropeptide-related proteins, such as FMRFa, AstC, their receptors and the enzymes involved in neuropeptide processing. A functional role of these candidates in nociception could be explored further by complementation of *C. elegans* lost sensory functions by the *O. vulgaris* orthologous gene. Overall, we highlighted an intersecting bioinformatic model hopping approach that facilitates understanding of nociception in *O. vulgaris* and which has key relevance for an evolutionary perspective on the phenomenon of pain.

## Acknowledgments

We thank Prof Laura Bianchi (University of Miami, Miami, FL) for kindly providing the double mutant *acd-1/deg-1 (bz90/u38)* and Dr Xiaofei Bai (University of Florida, Gainesville, FL) for kindly sharing AG405 *pezo-1 (av143)* strain. The rest of the strains were provided either by the *Caenorhabditis* Genetic Center (CGC), funded by NIH Office of Research Infrastructure Programs (P40 OD010440) or the National Bioresource Project (NBRP), funded by the AMED (Japan Agency for Medical Research and Development).

## Author contributions

Conceptualization: E.M.P., L.H., V.O., G.F., J.D.; Methodology: E.M.P., L.H., V.O., J.D.; *In silico* analysis: E.M.P. P.I.; Experimental investigation: E.M.P.; Resources: L.H., V.O., J.D.; Writing-original draft: E.M.P., L.H., V.O., J.D.; Supervision: L.H., V.O., J.D.; Project administration: L.H., V.O., J.D.; Funding acquisition: L.H., V.O., G.F.

## Data and resource availability

All relevant data and details of resources can be found within the article and its supplementary information

