## Supplementary figures and images for "Identification of molecular nociceptors in *Octopus vulgaris* through functional characterisation in *Caenorhabditis elegans*"

### FigS1

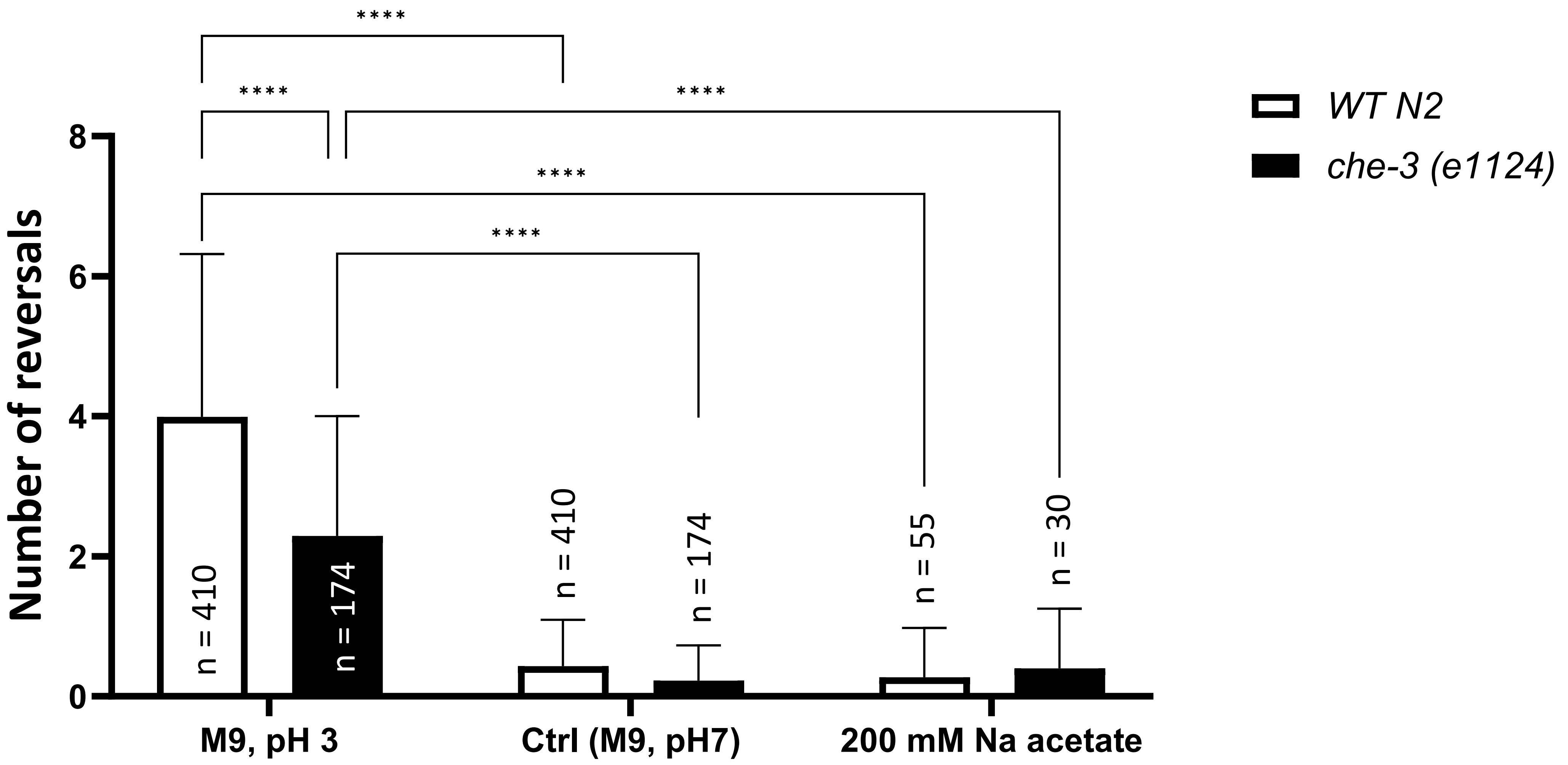
